# Dopamine alters the effect of brain stimulation on decision-making

**DOI:** 10.1101/2023.06.05.543812

**Authors:** Li-Ann Leow, Anjeli Marcos, Esteban Nielsen, David K Sewell, Tim Ballard, Paul E. Dux, Hannah L. Filmer

**Author notes:** Corresponding author: Li-Ann Leow. School of Psychology, McElwain Building, The University of Queensland, St Lucia, Australia.

## Abstract

Non-invasive brain stimulation techniques, such as transcranial direct current stimulation (tDCS), show promise in treating cognitive and behavioural impairments in clinical conditions. However, optimisation of such clinical applications requires a better understanding of how tDCS alters cognition and behaviour. Existing evidence implicates dopamine in the way tDCS alters brain activity and plasticity, however, there is as yet no causal evidence for a role of dopamine in tDCS effects on cognition and behaviour. Here, in a preregistered, double-blinded study, we examined how pharmacologically manipulating dopamine altered the effect of tDCS on the speed-accuracy trade-off, which taps ubiquitous strategic cognitive processes. Cathodal tDCS was delivered over the left prefrontal cortex and the superior medial frontal cortex before participants completed a dot-motion task, deciding the direction of moving dots under instructions to emphasize speed, accuracy, or both. We leveraged computational modelling to uncover how our manipulations altered latent decisional processes driving the speed-accuracy tradeoff. We show that dopamine in combination with tDCS (but not tDCS alone, nor dopamine alone) not only impaired decision accuracy, but also impaired discriminability, which suggests that these manipulations altered the encoding or representation of discriminative evidence. This is, to the best of our knowledge, the first direct evidence implicating dopamine in the way tDCS affects cognition and behaviour.

**Significance statement:** Transcranial direct current stimulation (tDCS) can improve cognitive and behavioural impairments in clinical conditions, however better understanding of its mechanisms is required to optimise future clinical applications. Here, using a pharmacological approach to manipulate brain dopamine levels in healthy adults, we demonstrate a role for dopamine in the effects of tDCS in the speed-accuracy trade-off, a strategic cognitive process ubiquitous in many contexts. In doing so, we provide direct evidence implicating dopamine in the way tDCS affects cognition and behaviour.

---

Transcranial electrical stimulation (tES) is low-cost, portable, has limited side effects, and is showing substantial promise across a range of applications, for example, in treating depression, stroke, and schizophrenia (Fregni *et al*., 2021). However, to fully realise the benefits of tES, a better understanding of *how* it affects brain function is necessary. Indeed, elucidating the mechanisms of tES is a prerequisite to optimise and more precisely target intervention protocols for better treatment efficacy.

A prominent form of tES is transcranial direct current situation (tDCS), where weak (0.5 to 4mA) currents are applied to the cortex via anodal and a cathodal electrodes placed on the scalp over brain area(s) of interest. Early conceptualisations attributed tDCS effects to changes in neural excitability: anodal stimulation shifted resting membrane potentials towards depolarisation, and cathodal stimulation towards hyperpolarisation (Bindman *et al*., 1962). However, recent in-vitro and in-vivo studies have painted a more complex picture.

Despite no changes to neural firing rates (Krause *et al*., 2017), tDCS can change the *timing* of neural firing (Reato *et al*., 2010), and alter synaptic plasticity (Márquez-Ruiz *et al*., 2012), even in regions distal to the electrodes (Rohan *et al*., 2015; Krause *et al*., 2023). Indeed, stimulation not only alters functioning of areas under the electrodes (Bachtiar *et al*., 2015), but can also impact activity and neurochemistry of functionally connected, remote regions. Notably, stimulating prefrontal cortex is associated with dopamine release in the basal ganglia (Fonteneau *et al*., 2018; Fukai *et al*., 2019; Bunai *et al*., 2021). In a similar vein, functional magnetic resonance imaging (Chib *et al*., 2013; Meyer *et al*., 2019) and direct electric field recording studies (Chhatbar *et al*., 2018) have shown that tDCS applied to the frontal cortex alters activity in midbrain basal ganglia circuits, and changes functional connectivity in the corticostriatal circuits (Polania *et al*., 2011); areas where dopamine orchestrates function (Calabresi *et al*., 1997).

The involvement of dopamine in the effects of tDCS is perhaps unsurprising given it’s importance in neural processes driving plasticity (Calabresi *et al*., 2007), motivation (Salamone & Correa, 2012), and cognition (Ott & Nieder, 2019). The bulk of evidence implicating dopamine in tDCS effects comes from pharmacological manipulations of dopamine which dramatically alters effects of tDCS on motor cortex plasticity (Nitsche *et al*., 2006; Monte-Silva *et al*., 2009; Monte-Silva *et al*., 2010; Fresnoza *et al*., 2014a; Ghanavati *et al*., 2022). For example, Levodopa, a dopamine precursor, increases the persistence of stimulation-induced neuroplastic aftereffects in the motor cortex by 20-fold (Kuo et al., 2008). Comparatively little is known about the role of dopamine in stimulation-induced changes in high-level cognition and behaviour. Whilst some studies suggest that markers of dopamine processing (e.g., dopamine genotype) partly predict responsivity to tDCS (e.g., Witte *et al*., 2012), evidence is mixed (Jongkees *et al*., 2019) and largely correlational. This is an important knowledge gap, as improved cognition and behaviour is often a core treatment goal in clinical applications of brain stimulation (Chase *et al*., 2020).

Here, we test the proposal that changes in brain dopamine function causally influence how tDCS modulates cognitive processes. To this end, we examined how dopamine perturbations alter the effect of tDCS on a well-studied phenomenon: the speed-accuracy trade-off (SAT). The SAT reflects strategic cognitive processes ubiquitous to virtually all task contexts that involves cortico–basal ganglia circuits (Lo & Wang, 2006; Forstmann *et al*., 2008; Van Veen *et al*., 2008; Bogacz *et al*., 2010), and is sensitive to tDCS (e.g., Filmer *et al*., 2021; Filmer *et al*., 2022). Here, we used our previously established stimulation and behavioural protocols (Filmer *et al*., 2021) where we deliver cathodal tDCS over the left prefrontal cortex and superior medial frontal cortex (SFMC) to examine how increasing brain dopamine availability, via the dopamine precursor Levodopa, impacts tDCS efficacy to modulate the SAT. Importantly, we leveraged evidence-accumulation modelling to uncover how these manipulations altered latent decisional processes driving the strategic cognitive operations underlying the SAT. Such approaches are not only more sensitive than typical behavioural measures of reaction time or accuracy (Stafford *et al*., 2020), but also enable deeper inferences about *how* decision-making behavioura are altered. We hypothesised that boosting dopamine availability would modulate the influence of stimulation on the speed-accuracy trade-off.

## Method Design

We employed a mixed factorial design, with between-subjects factors of Stimulation Site (PFC, SMFC) and drug (Levodopa, Placebo) and within-subject factors of stimulation (cathodal tDCS, sham tDCS) and response strategy instruction (speed emphasis, accuracy emphasis, or balance speed and accuracy equally).

Participants were randomly assigned to one of the four conditions (Levodopa PFC, Levodopa SMFC, Placebo PFC, PlaceboSMFC). Each participant took part in two sessions where they completed sham tDCS and cathodal tDCS, with session order counter-balanced across participants. In both sessions, after tDCS, participants completed a random dot-motion discrimination task where they were instructed to either emphasize speed, emphasize accuracy, or balance between speed and accuracy. The study was preregistered on the Open Science Framework, https://osf.io/gz5h9/.

### Participants

A total of 62 right-handed young adults (age: 18-35 years old, 24 males) with no known neurological/psychiatric conditions and no contraindications to brain stimulation or the drug manipulations participated in the study. The study was approved by the Human Research Ethics Committee at The University of Queensland and conformed to the Declaration of Helsinki. All participants provided written informed consent. Participants either received course credit at a rate of 1 course credit per hour or reimbursed with cash or gift cards, at a rate of $20/hour.

Sample size was determined via a Bayesian sampling rule, where we recruited participants until a BF_10_ > 3 or BF_01_ > 3 was established for the critical hypothesis test for the Drug x Stimulation interaction for any one of the four dependent variables (described in statistical analyses section), providing evidence for the alternative or null hypothesis, or until we collected 30 complete datasets for each condition, whichever was sooner. We achieved BF_10_ >3 for the critical hypothesis test at our sample of 62 participants.

### Procedure

All participants (regardless of whether they were assigned to the Levodopa condition or the placebo condition) completed two sessions, where they were subjected to cathodal tDCS or sham tDCS in each session (order counterbalanced). Figure 1A shows an overview of the session protocol. After initial blood pressure and mood assessments, participants received either placebo (vitamin) or levodopa (Madopar 125: 100 mg levodopa and 25 mg benserazide hydrochloride), crushed and dispersed in orange juice. An experimenter uninvolved in data collection prepared the solution in order to double-blind the study. Participants then completed the Barratt Impulsivity Scale (BIS), and two assessments of working memory (the operation span and memory updating task, order counterbalanced). These measurements were taken because individual differences in impulsivity and working memory capacity have been previously suggested to partly account for individual differences in dopamine synthesis capacity (Cools *et al*., 2008; Buckholtz *et al*., 2010). This was followed by a second blood pressure and mood rating assessment, and then electrode placement. After electrode placement, participants completed the practice and the thresholding phase of the dot-motion discrimination task. Stimulation commenced after the practice and thresholding phase, approximately 40 minutes after drug ingestion, within the window of peak plasma dopamine availability after Madopar consumption (Contin & Martinelli, 2010). After stimulation, participants completed the main task. Upon task completion, participants and experimenters completed questionnaires assessing blinding efficacy. A final blood pressure and mood rating assessment was also completed.

**Figure 1.**
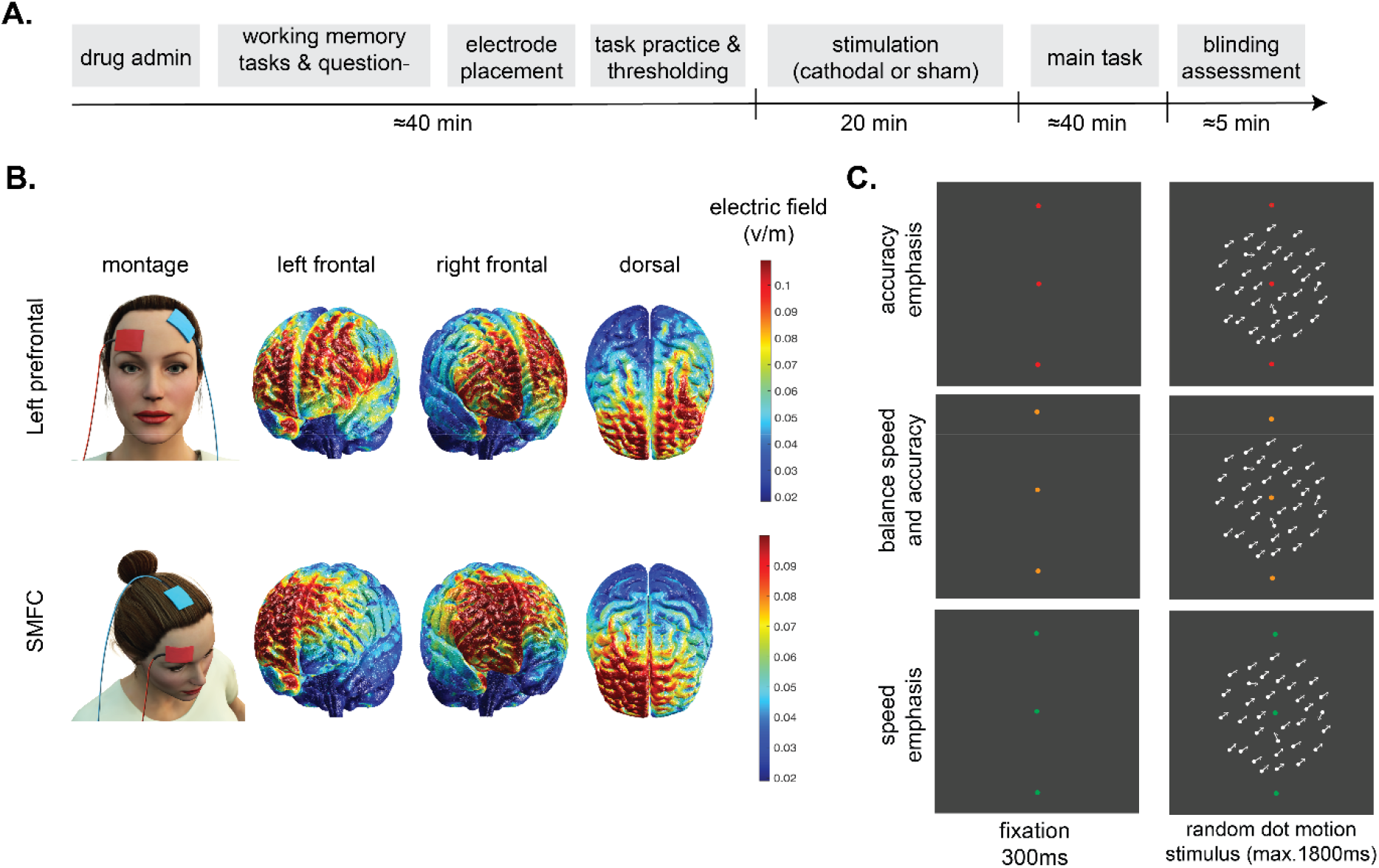
A. Session timeline, where each participant completed a sham stimulation session and a true stimulation session (order counterbalanced across participants). Participants received the same drug type (levodopa or placebo) across both sessions. B. Electrode montages and induced current modelled via ROAST (Huang et al., 2018) for the left prefrontal cortex stimulation site and the SMFC stimulation site. C. Trial outline for each response strategy type.

**Figure 2.**
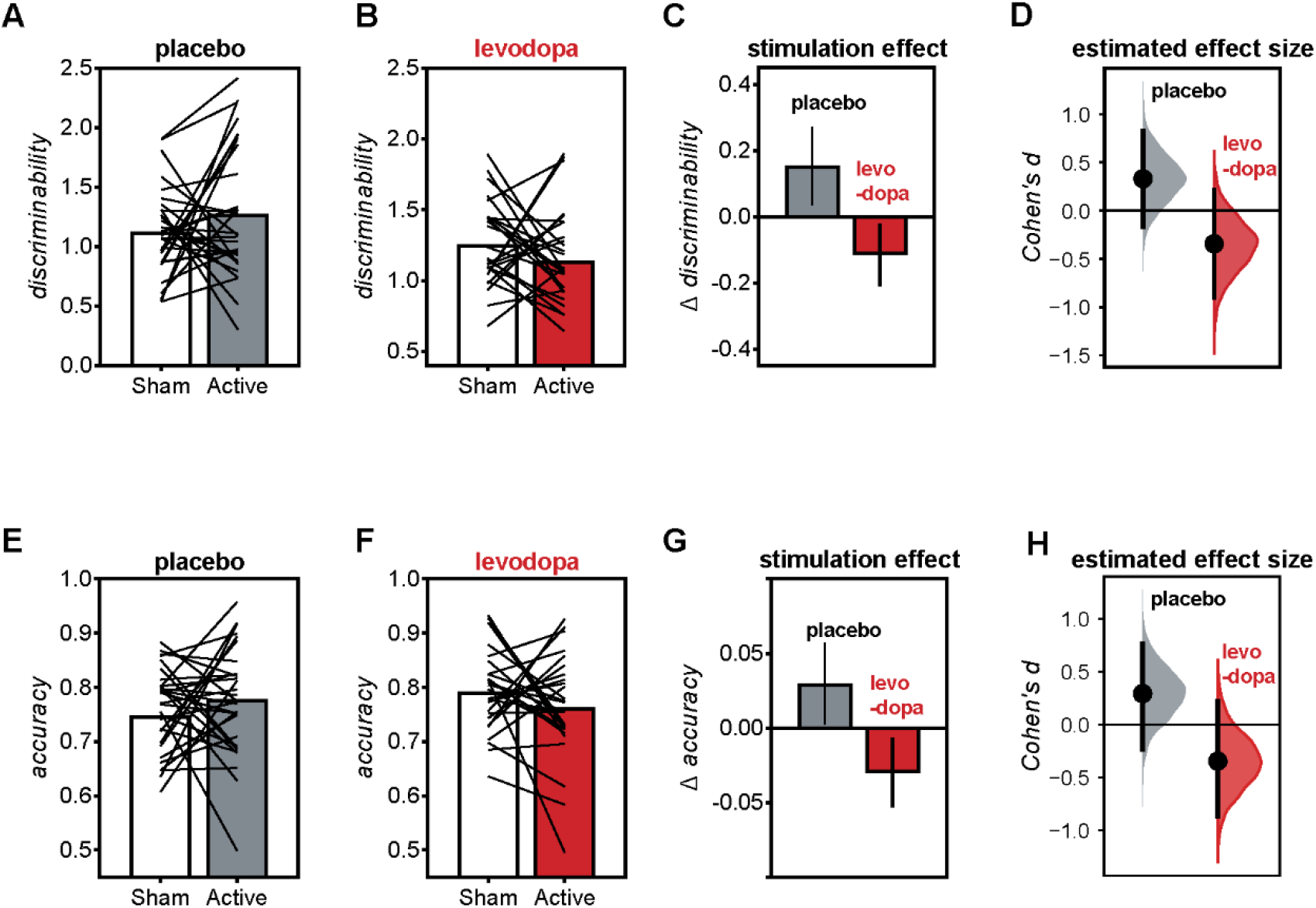
Mean discriminability (top panels) and mean accuracy (bottom panels) averaged across all response strategy conditions, under sham and active stimulation conditions, for participants who received either levodopa (B&E) or placebo (A&D). Effect of stimulation is shown in C (mean+/-SEM for change in discriminability from sham to cathodal stimulation) and F (mean +/-SEM for change in accuracy from sham to cathodal stimulation). Cumming’s estimation plots for effect size estimates (cohen’s d) for the effect of stimulation for placebo and levodopa for discriminability (D) and accuracy (H).

### Task

A random dot motion task was used to investigate decision making, similar to that employed previously (Filmer *et al*., 2021). Each trial started with a fixation dot, shown for 300 ms, followed by a random dot display (shown for 1800 ms) of 200 white dots (RGB: 255 255 255 background, presented within a circular area with a diameter of 3.4° visual angle and moving at a rate of 0.46° visual angle/second), against a black (RGB: 0 0 0) background. The random dot display contained a subset of dots (75%) that moved coherently either towards the upper left or right of the screen, while the rest of the dots moved randomly. The subset of coherent dots was determined on a frame-by-frame basis. Participants were instructed to discriminate the direction of coherent motion of the dots, indicating leftwards by pressing the “<“ key with their right index finger, and rightwards by pressing the “>“ key with their right middle finger, using an Apple Keyboard (A1243) to record keypresses. Stimuli were presented via a 2014 Mac mini (OS version: High Sierra) connected to a 24” VG248 Monitor (1920 × 1080 resolution; Intel Iris 1536 MB; 100 Hz refresh rate). The task was programmed in MATLAB (2016b) using the Psychtoolbox extension version 3.0.8 (Brainard, 1997).

At the start of each session, participants completed a short practice block (20 trials) with feedback on discrimination accuracy, followed by a thresholding procedure. The thresholding procedure determined the 75% correct direction discrimination via a QUEST staircase that varied the degree of motion offset from the vertex. Three independent QUEST staircases were used in this thresholding phase (Watson & Pelli, 1983), where the outcome of 90 trials was averaged to provide the threshold value for the main task. The staircase result was run for 20 more trials to give an estimate of the thresholding success. The thresholding procedure was rerun if performance on these trials was < 60% or > 90%.

After thresholding, participants practiced using the three different response strategies (speed emphasis, accuracy emphasis, or balanced between the two). The strategy instructions were blocked, with participants completing one block per strategy type (order randomised across subjects) for six trials per block. Figure 3 depicts the colour of the fixation dot at the start of the trial, which indicated the strategy to be used in a given block: green for prioritising speed (RGB: 0 255 0), orange for balancing speed and accuracy (RGB: 255 165 0), and red for prioritising accuracy (RGB: 255 0 0). Participants were instructed that speed and accuracy were both important and the fixation cues should prompt a shift in emphasis. Next, stimulation was administered (see below). Within 2 minutes of stimulation cessation, participants completed the main task, which consisted of 36 blocks (12 per response strategy). The block types were pseudo-randomly intermixed, controlling for the direction and distance of response strategy shifts, and ensuring the different block types occurred equally often across every 6 blocks of the experiment. The length of each block was an average of 25 trials (minimum of 20, maximum of 30, with a standard deviation of 3; approx. 300 trials per response strategy per session). The blocks lasted an average of 32 seconds, with a 30 second break given every 6 blocks. The main task lasted approx. 40 minutes.

**Figure 3.**
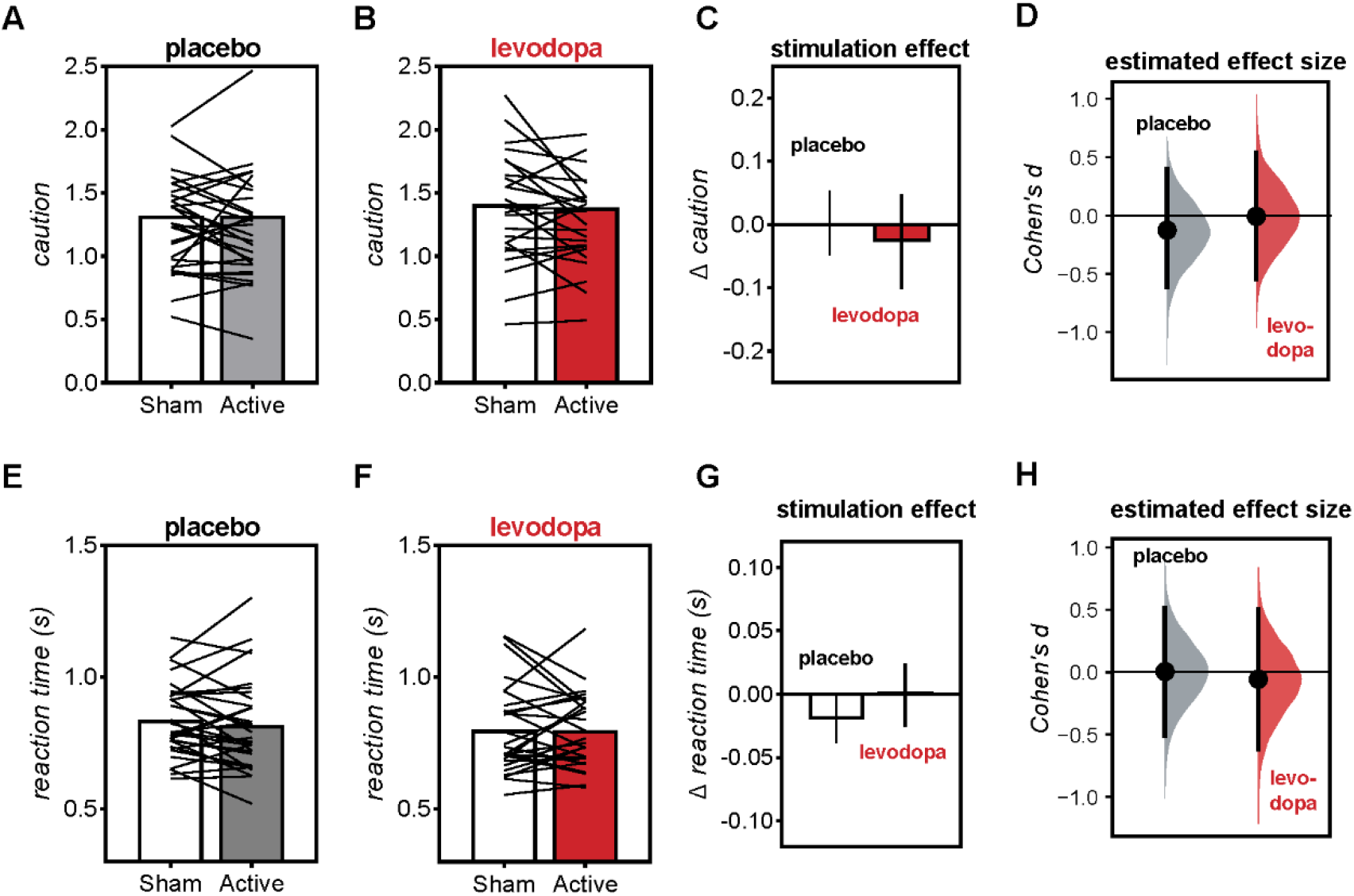
Mean response caution (top panels) and mean reaction time (bottom panels) averaged across all response strategy conditions for participants who received either levodopa or placebo. Mean+/-SEM for change in discriminability from sham to cathodal stimulation (C). Mean +/-SEM for change in discriminability from sham to cathodal stimulation (F). Cumming’s estimation plots for the effect size (cohen’s d) of stimulation for placebo and levodopa for discriminability (D) and accuracy (H).

### Stimulation

Participants received cathodal or sham stimulation to the left PFC or the SMFC (see Figure 1B). Electrode placement was determined via the EEG 10-20 system (Jasper, 1958): for the left PFC condition, the target electrode was placed 1 cm posterior to F3. For the SMFC condition, the target electrode was placed 1cm posterior to the Fz site. For both conditions, the reference electrode was placed over the contralateral supraorbital region (the area above the right brow ridge). Cathodal stimulation lasted for 20 minutes (30 second ramp up, 19 minutes constant stimulation at 0.7mA, and 30 seconds ramp down). For sham stimulation, the ramp-up/ramp-down parameters were the same, but during the 19 minute period, only 15 s of stimulation was delivered, and then small testing pulses were delivered to provide an active measure of impedance (to facilitate experimenter blinding) and to provide ongoing sensations to the participant. Stimulation was double-blinded using stimulation codes provided by a team member not involved in data collection. Stimulation was delivered via a NeuroConn stimulator with two 5 × 5 cm electrodes. Participants were instructed to sit quietly and keep their eyes open during stimulation.

### Additional Measures

Before electrode placement, we made additional assessments, intended to be entered as covariates in our analyses. We measured (1) trait impulsivity, measured via the BIS-11, a 30-item questionnaire with the following factors: attentional (attention and cognitive instability), motor (motor and perseverance), and non-planning (self-control and cognitive complexity) and (2) working memory, measured via the memory updating task and the operation span task from a working memory test battery (Lewandowsky *et al*., 2010), order counterbalanced for each session. We took these additional assessments because, at the time of planning the study, previous evidence suggested associations between trait impulsivity scores and working memory with dopamine synthesis capacity (Cools *et al*., 2008; Buckholtz *et al*., 2010), and impulsivity and working memory scores can partly predict the effect of dopamine agents on learning (van der Schaaf *et al*., 2014; Sharp *et al*., 2016; Froböse *et al*., 2018; Fallon *et al*., 2019). However, recent evidence using larger sample sizes has demonstrated an *absence* of a relationship between striatal dopamine synthesis capacity and working memory capacity, and between striatal dopamine synthesis capacity and impulsivity (van den Bosch *et al*., 2022).

Consistent with this recent evidence (van den Bosch *et al*., 2022), inclusion of the covariates failed to change any of the conclusions made in our analyses. Furthermore, across all analyses, we found moderate-strong evidence to exclude the covariates in the analyses. Thus, for parsimony, we did not include these covariates in our final analysis.

### Blinding

Blinding efficacy for both the experimenter and the participant was assessed at the end of each session, by asking both the experimenter and the participant to complete online forms which ask whether they thought the participant received (1) placebo or Madopar, and (2) true tDCS or sham tDCS.

### Data analysis

Data exclusion criteria: To minimize data loss, we made a small deviation from our preregistered exclusion criteria. Instead of excluding participants who showed floor (< 60%) or ceiling (> 90%) performance for any block of the three strategy conditions, we now exclude participants with floor (< 60%) performance, as indicated by mean accuracy averaged across all strategy conditions in each test session. According to this criteria, two subjects from the PFC group were removed, and three subjects from the SMFC group were removed from the analyses.

## Computational modelling

We used the linear ballistic accumulator (LBA) model within a hierarchical Bayesian framework (Brown & Heathcote, 2008) to quantify the latent components of the decision process. For each accumulator, the rate at which evidence accumulates (the drift rate) is drawn from a normal distribution with mean v and standard deviation sv. Evidence accumulates until the evidence for one alternative breaches a threshold b, at which point that alternative is selected and a response is made. As is common practice (e.g., (Brown & Heathcote, 2008), we express the threshold as the difference between the raw threshold and the maximum starting point (denoted B, where B = b – A). The LBA also includes a nondecision time parameter (denoted t0) that captures the component of the reaction time (RT) attributable to processes other than the decision process (e.g., encoding the stimulus and manually executing the response).

The model was parameterized such that the mean rate of evidence accumulation for the correct response alternative could vary across experimental conditions, whereas the mean rate for the incorrect response alternative was fixed across experimental conditions. This was done to enable a more reliable measure of discriminability and caution (see (Evans, 2020)). **Discriminability** was indexed by the difference between the mean rate of evidence accumulation for the response alternative that was correct and the mean rate for the response alternative that was incorrect (this difference is indexed by the *vdiff* parameter). **Response caution** was quantified via the threshold parameter (B), which was also allowed to vary across conditions. Nondecision time (t0) and the maximum starting point of evidence (A) were fixed across conditions. The standard deviation of the rate of evidence accumulation was fixed to one for both accumulators in all conditions.

The model also assumed that the mean rate of evidence accumulation for the correct response and threshold could vary as a function of stimulation site and drug. As these factors were manipulated between participants, their effects were modelled by assuming that the rate for the correct response and the threshold for the different participant groups were drawn from different population distributions. The priors on the population parameters were based on the recommendations of Gronau *et al*. (2020). The participant-level parameters were modelled using truncated normal distributions. Non-decision time (t0) was bounded at 0.1 and 1. All other parameters had a lower bound of 0 and no upper bound.

### Statistical Analyses

We considered four dependent variables: response accuracy, reaction times for correct trials, response caution, and response discriminability.

To test the effect of stimulation, drug manipulation, and stimulation site on performance across different response strategy conditions, we ran Stimulation (cathodal, sham) x Drug (levodopa, placebo) x Strategy (speed emphasis, accuracy emphasis, balanced) x Site (SMFC, PFC) Bayesian ANOVAs on all four dependent variables.

BF values of 1–3 were interpreted as weak, 3–10 as moderate, and > 10 as strong evidence for either the test hypothesis/variable inclusion (BF_10_ or BF_incl_) or for the null hypothesis/variable exclusion (BF_01_ or BF_excl_). BF values ∼1 were interpreted as providing no evidential value. JASP 0.16.2 was used for all statistical analyses.

We also plotted Cumming estimation plots for Cohen’s d for effects of stimulation (i.e., comparing sham stimulation versus cathodal stimulation) for all four dependent variables, separately for the Levodopa and placebo groups, using DABEST (‘data analysis with bootstrap-coupled estimation’) via the web application at https://www.estimationstats.com (Ho et al., 2019).

## Results Manipulation checks

Consistent with previous findings (Filmer *et al*., 2021), the response strategy instructions had a clear effect on reaction time and response caution, as shown by a main effect of instruction on reaction time (BF_incl_ = 21,072) and response caution (BF_incl_ = 9.382e+9). Also consistent with previous work, instructions had no effect for accuracy (BF_excl_ = 4.622) or discriminability (BF_excl_ = 9.605). Instructions emphasising speed shortened reaction times and reduced response caution compared to instructions emphasised balancing between speed and accuracy (reaction time: BF_10_ = 1.011e+8, response caution: BF_10_ = 2.498 e+6), and compared to instructions emphasizing accuracy (reaction time: BF_10_ = 1.463e+9, response caution: BF_10_ = 1.260 e+7). In addition, instructions emphasizing accuracy slowed reaction times and increased response caution compared to the balanced emphasis condition (reaction time: BF_10_ = 1012.354, response caution: BF_10_ = 317.327).

### Levodopa alters stimulation effects on accuracy and discriminability

Levodopa altered the effect of stimulation for response accuracy and discriminability, as shown by strong evidence for a Drug x Stimulation interaction on both these dependent variables (accuracy BF_incl_ = 33.316; discriminability BF_incl_ = 34.098). Follow-up Stimulation x Strategy ANOVAs were run separately for the levodopa and the placebo groups.

For the levodopa group, there was moderate evidence for the main effect of stimulation for accuracy (BF_incl_ = 5.501) and weak evidence for the main effect of stimulation for discriminability (BF_incl_ = 2.811), where cathodal tDCS reduced response accuracy compared to sham (cathodal: M = 0.763 [95%CI: 0.729, 0.797]; sham: M = 0.792 [95%CI: 0.758, 0.827]), and reduced discriminability compared to sham (sham: M = 1.255 [95%CI: 1.121, 1.388]; cathodal M = 1.144 [95%CI: 1.010, 1.277]).

For the placebo group, there was indeterminate evidence for the main effect of stimulation on accuracy (BF_excl_ = 0.644), and moderate evidence for excluding the Stimulation x Strategy interaction (BF_excl_ = 9.158).

There was weak evidence for the main effect of stimulation on discriminability (main effect of stimulation, BF_incl_ = 2.626), mean discriminability was numerically higher with cathodal stimulation (M = 1.277 [95%CI:1.107, 1.447]) than with sham stimulation (M = 1.122 [95%CI:0.952, 1.292]), consistent with previous results (Filmer *et al*., 2021).

### Levodopa does not alter effect of stimulation on reaction times or response caution

We found evidence that Levodopa did not alter the effect of stimulation on reaction times and response caution, as we saw moderate evidence for excluding the Stimulation x Drug interaction from the model, both for response caution (BF_excl_ = 5.537), and reaction times (BF_excl_ = 3.588), and moderate evidence for excluding the Stimulation x Drug x Strategy interaction, both for response caution (BF_excl_ = 7.966) and reaction times (BF_excl_ = 9.640).

### Effect of stimulation site

Whilst we chose to use two different stimulation sites, in keeping with our previous work, our stopping rule was based on pooled data from both stimulation sites reaching BF>3. Thus, our sample size was not optimised to test for either the effect of stimulation (under the placebo conditions), or the effects of stimulation site. The sample sizes that we used for each stimulation site and each stimulation condition under no-drug conditions were therefore smaller than that used in previous studies focussing on effects of stimulation on the speed-accuracy tradeoff (Filmer *et al*., 2022). We did not find an effect of stimulation site, as shown by moderate evidence for excluding the Site x Stimulation interactions from the models on accuracy (BF_excl_ = 3.769), reaction times (BF_excl_ = 4.115), discriminability (BF_excl_ = 6.110) and response caution (BF_excl_ = 5.531).

### Control analyses

To evaluate whether levodopa had non-specific effects on decision-making (in isolation from the effect of stimulation), we ran Strategy x Drug ANOVAs on the sham session data. Overall, in the absence of stimulation, there was weak-to-moderate evidence for excluding the effect of Levodopa. In the absence of stimulation, Levodopa did not have prominent effects on reaction times, accuracy, response caution, or discriminability regardless of speed/accuracy emphasis, as shown by evidence for excluding the main effect of drug [discriminability: BF_excl_ = 1.191, accuracy: BF_excl_ = 0.790; response caution; BF_excl_ = 6.813, reaction times (BF_excl_ = 4.532)], and weak-moderate evidence for excluding the drug x instruction interaction [discriminability: BF_excl_ =1.191, accuracy: BF_excl_ = 12.668; response caution; BF_excl_ = 6.813, reaction times (BF_excl_ = 4.532)].

### Blinding Participant blinding

Overall, participants did not appear to be above chance at correctly guessing the stimulation and drug conditions. Specifically, Bayesian binomial tests showed that, in session 1, there was weak evidence for participants being above chance at correctly guessing the stimulation condition (BF_10_ = 1.353), and moderate evidence that participants were not above chance at correctly guessing and the drug manipulation (BF_01_ = 3.591). In session 2, there was weak evidence for the participants being not above chance at correctly guessing the stimulation condition (BF_01_ = 1.952) and the drug condition ((BF_01_ = 1.952).

### Experimenter blinding

Overall, experimenters did not appear to be above chance at correctly guessing the stimulation and drug conditions. Specifically, in session 1, there was strong evidence for experimenters being not above chance at correctly guessing the stimulation condition (BF_01_ = 11.439) and weak evidence for the experimenter being above chance at correctly guessing the drug manipulation (BF_01_ =1.785). In session 2, there was moderate evidence for experimenters being not above chance at correctly guessing the stimulation condition (BF_01_ = 7.060) and strong evidence for the experimenter being not above chance at correctly guessing the drug manipulation (BF_0+_ = 13.041).

### Control measurements

Pharmacological manipulations of dopamine can occasionally elicit undesired side effects such as nausea, resulting in some participants withdrawing from the study (for example, see Chen *et al*., 2020), and can also change mood state (Vo *et al*., 2018). None of our participants reported nausea. We did not find evidence demonstrating that levodopa altered our participants’ blood pressure, as Time (before drug administration, 1 hour after drug administration, 2 hours after drug administration) x Drug ANOVAs yielded weak-to-strong evidence for the null hypothesis for Drug x Time interactions for blood pressure (diastole: BF_excl_ = 13.985, systole: BF_excl_ = 2.361), mood (BF_excl_ =7.473) and heart rate (BF_excl_ = 15.333).

## Discussion

In a pre-registered, double-blind study, we tested the proposal that transcranial direct current stimulation alters behaviour, partly by altering dopamine function. To this end, we examined how increasing dopamine availability in neurotypical young adults causally altered the effect of tDCS on the speed-accuracy trade-off. We leveraged behavioural and computational modelling protocols known to be sensitive to the effect of non-invasive brain stimulation on the speed-accuracy trade-off (Georgiev *et al*., 2016; Berkay *et al*., 2018; Filmer *et al*., 2021; Filmer *et al*., 2022). Levodopa altered the effect of stimulation on response accuracy and response discriminability, and effects were selective to the active but not the sham stimulation conditions. The absence of an effect of Levodopa in the sham stimulation condition argues against the possibility that increasing dopamine availability via Levodopa had non-specific effects on performance. The finding that Levodopa modulates the effect of brain stimulation on response strategy during decision-making is, to the best of our knowledge, the first causal evidence for a role of dopamine in how frontal tDCS alters *behaviour* in neurotypical humans. Further, our modelling approach also allows us to demonstrate that combining tDCS and Levodopa changed decision-making behaviour by altering process(es) responsible for encoding or representing discriminative evidence.

Previous work directly implicating dopamine in the effects of non-invasive brain stimulation have focussed on the neurophysiological effects of motor cortex stimulation, by detailing how different dopamine perturbations changes the way stimulation affects markers of motor cortex plasticity (Kuo *et al*., 2008; Monte-Silva *et al*., 2009; Monte-Silva *et al*., 2010; Fresnoza *et al*., 2014a; Fresnoza *et al*., 2014b). For example, medium doses of Levodopa (100mg) combined with cathodal tDCS of the motor cortex resulted in a 20-fold increase in the persistence of the suppression of motor evoked potentials after cathodal tDCS (Kuo *et al*., 2008). Such findings do not, however, allow us to make direct inferences about the way tDCS alters behaviour and cognitive strategy adjustment. Furthermore, findings on the motor cortex might not generalise to other brain regions such as the prefrontal cortex, as the motor cortex possesses distinct cytoarchitecture (Sanes & Donoghue, 2000), is part of a distinct cortico-striatal pathway (Alexander *et al*., 1986), and thus shows distinct patterns of responsivity to brain stimulation compared to the frontal cortex (Jamil *et al*., 2017).

By showing that the dopamine precursor Levodopa altered effects of brain stimulation on decision-making, we provided direct evidence for a role of dopamine in the mechanism of how brain stimulation alters cognition and behaviour. To date, evidence for the role of dopamine in stimulation effects on behaviour has been correlational. For example, genetic polymorphisms which partly account for individual differences in dopamine function has been associated with sensitivity to non-invasive brain stimulation (Witte *et al*., 2012; Plewnia *et al*., 2013; Nieratschker *et al*., 2015); although effects are inconsistent, (c.f., Jongkees *et al*., 2019). Similarly, work in humans and in animals shows that frontal cortex tDCS is associated with altered activity in the subcortical networks (Keeser *et al*., 2011; Weber *et al*., 2014; Hunter *et al*., 2015), as well as with midbrain dopamine release (Tanaka *et al*., 2013; Fonteneau *et al*., 2018; Bunai *et al*., 2021). One PET study showed only a weak correlation between post-stimulation dopamine release with reaction time variability (Fukai *et al*., 2019). By demonstrating that perturbing brain dopamine levels alters the effects of tDCS on behaviour, the current study provides direct evidence for the role of dopamine in the way tDCS affects cognition and behaviour, and is an important extension to the growing body of evidence that implicates dopamine in the effects of brain stimulation.

*How* might dopamine have altered the effects of frontal cortex tDCS on behaviour? One clue comes from studies of brain stimulation and neurochemistry. Brain stimulation can alter the excitation/inhibition balance between excitatory glutamate and inhibitory GABA function in the stimulated brain area (Kim *et al*., 2014; Heimrath *et al*., 2020), which may partly explain why the efficacy of stimulation depends partly on baseline GABA/glutamate levels (Filmer *et al*., 2019). Intriguingly, recent work has shown that stimulation is associated with reduced GABA levels not only in the area under the electrode, but also in the ipsilateral striatum, which negatively correlated with post-stimulation levels of striatal dopamine release: the greater the stimulation-associated decrease in GABA, the larger was the post-stimulation dopamine release (Bunai *et al*., 2021). In other words, frontal cortex tDCS is associated with a disinhibitory effect on the midbrain, as well as with increased midbrain dopamine release (Bunai *et al*., 2021). Frontal cortex stimulation might thus alter the inhibitory tone in the GABAergic pathways in the fronto-striatal loops. Whilst we do not yet fully understand the functional significance of these stimulation-induced changes in striatal GABA and dopamine levels, we speculate that here, combining protocols (i.e., exogenous dopamine via Levodopa and frontal cortex stimulation) which both increase dopamine release disrupted the typically optimal dopamine levels in the brain areas contributing to decision-making performance, ultimately reducing response accuracy and discriminability during decision-making. Such findings are in line with extant theories of an inverted-U shaped relationship between dopamine levels in the cortico-striatal circuits and cognitive function (e.g.,Cools & D’Esposito, 2011), which resulted from a large body of findings showing that increasing dopamine levels beyond optimal levels can impair cognition (e.g.,Arnsten *et al*., 1994; Mehta *et al*., 2001; Mehta *et al*., 2008; Gallant *et al*., 2016; Vo *et al*., 2016).

A feature of the current findings is the selectivity of the way in which dopamine perturbations altered stimulation effects: Levodopa only altered response accuracy and discriminability when combined with active stimulation: no effect was evident when Levodopa was combined with sham stimulation. The absence of an effect of Levodopa in the sham stimulation condition is consistent with previous findings showing evidence for no effect of dopamine drug manipulations on either response caution or discriminability in simple decision-making tasks (e.g., Winkel *et al*., 2012). Dopamine drug manipulations seem to only alter response caution or discriminability under specific circumstances: for example when decision-making was embedded within paradigms requiring proactive inhibition (Rawji *et al*., 2020), reinforcement learning (Chakroun *et al*., 2022; Mathar *et al*., 2022), or temporal discounting (Wagner *et al*., 2020), or only in clinical populations with impaired dopamine function and decision-making behaviour (Huang *et al*., 2015; Wagner *et al*., 2020). Studies which have found effects of dopamine drug manipulations on the speed-accuracy trade-off ((e.g., Pedersen *et al*., 2017; Beste *et al*., 2018; Bensmann *et al*., 2019) have used methylphenidate, a drug that works by blocking the re-uptake of dopamine and noradrenaline, resulting in increased dopamine levels in *both* frontal cortex (Claussen & Dafny, 2014) and the basal ganglia (Westbrook *et al*., 2019). Similar to our results which showed that combined levodopa and tDCS altered discriminability and accuracy without changing response threshold or reaction times, methylphenidate selectively altered discriminability and accuracy without changing response threshold or reaction times (Beste *et al*., 2018; Bensmann *et al*., 2019). Methylphenidate’s dual mode of action in both the frontal cortex and the striatum contrasts to other dopamine agents such as Levodopa which selectively act upon the striatum rather than the frontal cortex (Carey *et al*., 1995). One possibility is that concurrent changes in activity in both the frontal cortex as well as the striatum are required to alter decision-making, consistent with extant theories suggesting that coordinated activity in the corticostriatal pathways are required to drive decision-making behaviour (Wei *et al*., 2015). According to such theories, cortical areas representing different response alternatives change their firing rate as they appear to accumulate and integrate evidence for each alternative (Gold & Shadlen, 2007), and signalling across the cortico-striatal pathways drives the basal ganglia to disinhibit output nuclei, resulting in the selection and execution of the winning response alternative (Lo & Wang, 2006). We speculate that here, increasing dopamine availability in the striatum might have altered cortico-striatal signalling, priming the frontal cortex to the effects of cathodal tDCS, impairing the process of evidence accumulation and/or integration in frontal cortex areas, which ultimately impaired discriminability.

### Future directions

We only tested effects of tDCS on behaviour at a single dose of Levodopa. In the case of the motor cortex, it is clear that dopamine has dose-dependent effects on stimulation-induced plasticity: whilst medium doses of Levodopa augments the effects of stimulation (Kuo *et al*., 2008), smaller or larger doses of Levodopa reverses or removes the effect of stimulation (Monte-Silva *et al*., 2009; Monte-Silva *et al*., 2010; Fresnoza *et al*., 2014a; Fresnoza *et al*., 2014b). Whilst such inverted-U shaped relationships between the dose of brain dopamine manipulations and the post-stimulation aftereffect have been well-established for the motor cortex (Monte-Silva *et al*., 2009; Monte-Silva *et al*., 2010; Fresnoza *et al*., 2014a; Fresnoza *et al*., 2014b), we do not yet know if similar relationships exist between brain dopamine levels and frontal tDCS effects on behaviour and cognition. Future studies are needed to investigate the nature of the relationship between brain dopamine levels and stimulation effects on behaviour.

Exactly how dopamine might alter physiological effects of tDCS also remains incompletely understood. It is well established that tDCS-induced plasticity depends on activity at the N-methyl-d-aspartate (NMDA) receptors (Nitsche *et al*., 2003), which receives excitatory glutamate, leading to calcium influx required for changes in synaptic plasticity (Carey *et al*., 1995). One recent study has demonstrated that NMDA receptor antagonists can reverse the effect of dopamine agonists on stimulation-evoked neuroplastic effects in the motor cortex (Ghanavati *et al*., 2022), which is broadly consistent with a role of dopamine in long-term synaptic plasticity (Calabresi *et al*., 2007). Whilst we demonstrate a link between dopamine and tDCS effects on decision-making, it is clear that other neuromodulators such as serotonin and noradrenaline also play a prominent role in the effects of tDCS (Nitsche *et al*., 2009b; Batsikadze *et al*., 2013), which await clarification in future work. Better understanding of the neurochemical mechanisms how tES alters the brain and behaviour is essential for evidence-based optimisation of tES protocols in clinical conditions characterised by neurochemical disturbances, such as schizophrenia and Parkinson’s disease (Chase *et al*., 2020).

## Conclusion

Existing work implicated dopamine in tDCS-induced neuroplasticity (Kuo *et al*., 2008; Monte-Silva *et al*., 2009; Nitsche *et al*., 2009a; Nitsche *et al*., 2010; Fresnoza *et al*., 2014a; Fresnoza *et al*., 2014b) and demonstrated that tDCS induces midbrain release of dopamine (Fonteneau *et al*., 2018; Fukai *et al*., 2019; Bunai *et al*., 2021). However, until the present work, there was no direct evidence implicating dopamine in the effects of tDCS on higher-order cognition and behaviour. Here, leveraging a pharmacological approach to manipulate dopamine, we provide the first causal evidence for a role of dopamine in the way tDCS alters strategic cognitive operations underpinning the speed-accuracy tradeoff. The work will contribute to realising the full benefits of tES, by contributing to our understanding of *how* stimulation affects brain function, thus helping to elucidate the mechanisms of tES to optimise and more precisely target intervention protocols for better treatment efficacy in the future.

